# Training Experimentally Naive Seals for Vocal Learning Experiments

**DOI:** 10.1101/2024.08.27.609954

**Authors:** Diandra Duengen, Andrea Ravignani

**Affiliations:** Comparative Bioacoustics Research Group, Max Planck Institute for Psycholinguistics, Nijmegen, The Netherlands; Zoo Cleves (“Tiergarten Kleve”), 47533 Kleve, Germany; Center for Music in the Brain, Department of Clinical Medicine, Aarhus University & The Royal Academy of Music, Aarhus, Denmark; Department of Human Neurosciences, Sapienza University of Rome, Rome, Italy; Research Center of Neuroscience “CRiN-Daniel Bovet”, Sapienza University of Rome, Rome, Italy

**Keywords:** Animal training, vocal learning, vocal shaping, pinniped, Phoca vitulina

## Abstract

Harbor seals (*Phoca vitulina*) are a common zoo species that show a scientifically valuable propensity for vocal learning. Under human care, the seals can be trained to associate vocalizations with cues. This ability is termed vocal usage learning and is characterized by learning to use a vocalization in a specific context. Among mammals, seals are prime candidates to investigate vocal learning. Yet, only a handful of reports exist on harbor seal vocal learning abilities, and even fewer document how these were trained or tested. Here, we investigate how, and if, two experimentally naive harbor seals under human care can be trained to participate in scientific experiments. We describe the training and testing of two seals in two basic vocal learning experiments. We trained the animals to vocalize upon the presentation of discriminative stimuli (S_D_) through operant conditioning methods and tested their abilities to i) vocalize and refrain from vocalizing on two distinct S_D_’s, and ii) produce two different vocalizations upon the presentation of two different S_D_’s. Both seals learned the tasks: the first task was achieved within 118 trials (22 errors to criterion) and 220 trials (40 errors to criterion), the second task within 480 trials (158 errors to criterion) and 380 trials (94 errors to criterion), respectively. Our results confirm that harbor seals are capable of vocal usage learning and further suggest that associating individually distinct vocalizations with different S_D_’s may be more cognitively demanding than vocalizing and being silent on S_D_.

## Introduction

A single case study found that a male harbor seal (*Phoca vitulina*) can be trained to vocalize on a discriminative stimulus (S_D_) (Schusterman, 2008). In his review of vocal learning in pinnipeds, Schusterman (2008) for the first time described the fruitful training of vocal variability in a harbor seal. He brought one vocalization under stimulus control and selectively shaped it towards a novel ‘wa-wa’ sound (Goncharova et al., 2024; Schusterman, 2008). Thereby, his case study demonstrates vocal usage learning: the ability to use a vocalization, be it learned or innate, in a novel context as a result of experience (Janik & Slater, 2000; Jarvis, 2019). The report provides a valuable foundation to document vocal learning and shaping, potentially inspiring further research. A remarkable case of mammalian vocal production learning, plasticity in sound production, was a captive male harbor seal, who in the 1980’s showed untrained imitation of human speech (Duengen et al., 2023; Ralls et al., 1985). A later attempt to replicate vocal imitation in this species failed (Moore, 1996). Harbor seals are popular in zoos, amongst others for their comparatively easy husbandry and trainability. This, taken together with their propensity for vocal learning, makes them a good study organism. Beyond the contribution to research, the participation in (e.g., acoustic) experiments can enrich individuals, thereby improving animal welfare.

Under human care, vocal usage learning can be tested through operant conditioning experiments (Adret, 1993; Janik & Slater, 2000; Lattenkamp et al., 2018; Seyfarth & Cheney, 2010; Stoeger & Baotic, 2021; Vernes et al., 2021), in which animals are taught to vocalize upon the presentation of an arbitrary S_D_ (e.g., a hand sign, see Table 3.1). Vocal learning via operant conditioning was found in a closely related seal species, the gray seal (*Halichoerus grypus*; Shapiro et al., 2004; Stansbury et al., 2015). Dissecting vocal learning into its complex features – from “simple” vocalizing upon the presentation of an S_D_ to vocalizing and withholding vocalizations upon different S_D_’s, to producing different vocalizations upon other S_D_’s – requires a chain of trained behaviors (Janik & Slater, 2000; Shapiro et al., 2004; Vernes et al., 2021).

**Table 1:** Description of visual SD’s used to elicit the desired behaviors of the seals.

| Behavior | Seal | $S_D$ |
| --- | --- | --- |
| Vocalize | E & J | Hand next to the trainer's right ear |
| Silence | E & J | Stretched arm in front of the trainer's chest with open hand |
| Vocalization E1 | E | Hand next to the right ear |
| Vocalization E2 | E | Pointed finger rolling in 90-degree angle in front of the trainer |
| Vocalization J1 | J | Right hand and arm above the trainer's head |
| Vocalization J2 | J | Full hand in front of the trainer's mouth with a slight forward movement |

Building on published work in other species (Shapiro et al., 2004; Vergara & Barrett-Lennard, 2017), we describe the training and testing process to probe vocal learning in two harbor seals. Our approach combines classical training methods (e.g., operant conditioning through positive reinforcement, associative learning, shaping by successive approximations) and randomized trial orders (Gellermann, 1933). We prepared two experimentally naive harbor seals for a series of vocal learning experiments (Duengen et al., 2024) by training them to vocalize, withhold from vocalizing, shaping a novel vocalization type and bringing two different vocalizations under stimulus control.

## Material and Methods

### Setting

The seals were housed in a 230,000-liter freshwater pool in a 300 m^2^ enclosure at Zoo Cleves, Germany. Two of the six resident harbor seals were selected to participate in this study as they were vocally active individuals. Seal E was a 6-year-old female born at Zoo Osnabrück, Germany. Seal J was a 16-year-old male born at Zoo Duisburg, Germany. A third, vocally active seal dropped out of the experiment due to a severe eye condition. All seals were previously trained to participate in shows, which included behaviors such as object retrieval or flipper slapping. Shows did not take place during the period of the study. Training sessions consisted of general training, conducted by zoo staff, and research training, conducted by the experimenter. All training sessions took place in the main enclosure, where all seals resided, up to three times/week. Seals were tested separately, up to five times per week in the research enclosure, a partitioned area attached to the main enclosure. *Training* and *Testing* were conducted during visitor hours.

### Procedure and Training

Operant conditioning was facilitated through positive reinforcement, where a correct response was immediately followed by a bridge stimulus (here, a whistle) and fish as a primary reinforcer. Incorrect behavior resulted in a least-reinforcement-scenario (neutrality for ca. 3 seconds). Positive reinforcement was delivered through pre-cut pieces of mackerel (*Scomber scrombus*) and herring (*Clupea harengus*).

We distinguish between *Training* and *Testing* (see Fig. 1). Training sessions were conducted with all seals in the main enclosure, a phase in which multiple behaviors were trained. These sessions served to establish novel behaviors and maintain known behaviors. Establishing novel behaviors was needed for a series of vocal learning experiments (Duengen et al., 2024) and consisted of e.g., stationing and targeting behaviors, as well as “go into the water”, and “follow the trainer”. Much of this training time included habituating the animals to a novel research enclosure, where the experiments would later take place (see description below). Training novel behaviors was facilitated through capturing (e.g., “go into the water”), luring (e.g., “follow the trainer”), or shaping (“target”). In *Testing*, the seals participated in two dedicated tasks in a separate research enclosure. These tasks consisted of two different experiments, *Experiment 1: Vocalize vs. Silence* and *Experiment 2*: *Produce Different Vocalizations Upon Different S*_*D*_*’s*.

**Fig. 1:**
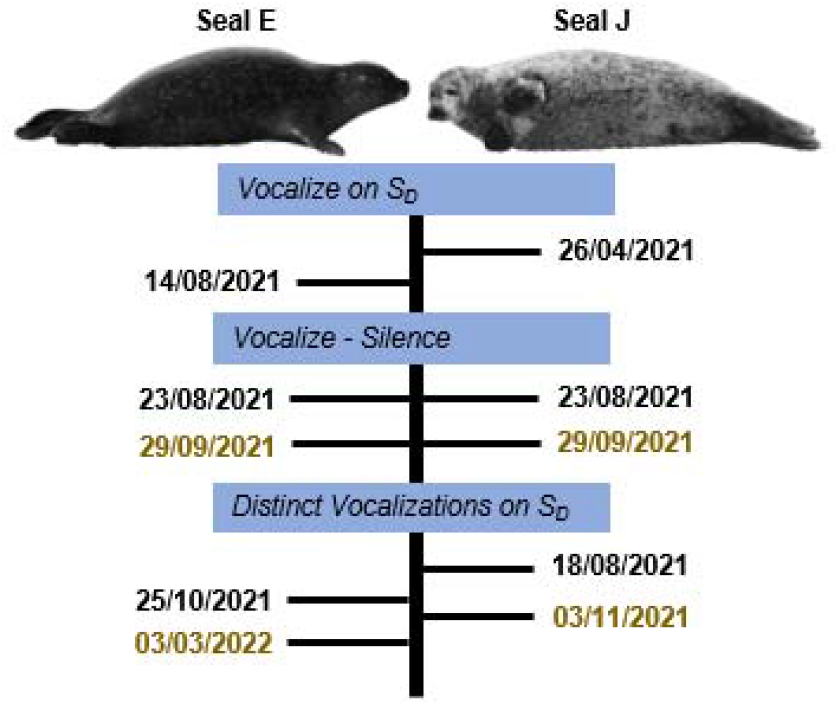
Illustration of the training (black) and testing (brown) timeline for seal E (left) and seal J (right).

#### Vocal Training

Seal E and Seal J would occasionally vocalize during general training sessions in April and August 2021. This behavior was captured by opportunistically reinforcing the vocal behavior with fish to increase its occurrence. Seal E offered one vocalization while Seal J offered two different vocalizations. Seal E’s second vocalization was shaped through successive approximations: The initial vocalization was rewarded whenever it aurally differed from the original vocalization and was produced with an opened muzzle. This led to the formation of a second type of vocalization (E2). When the seals produced vocalizations, we reinforced these with fish and presented a respective hand sign to facilitate the formation of an association between the vocal behavior and the S_D_ (see Table 1, Fig. 2).

**Fig. 2:**
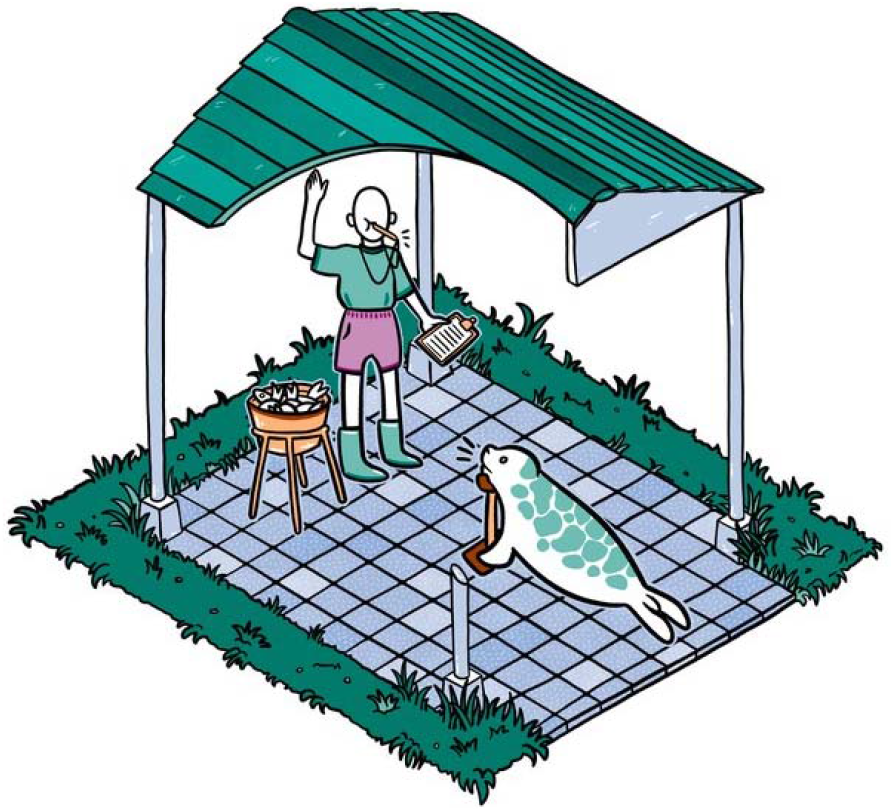
Illustration of the experimental Setup. The experimenter stands in front of the seal and asks for a pre-determined behavior (here: Vocalization J1, see Table 1). Responses were noted on the sheet, which contained the randomized order of behaviors (Gellermann, 1933; Jadoul et al., 2023b).

**Fig. 3:**
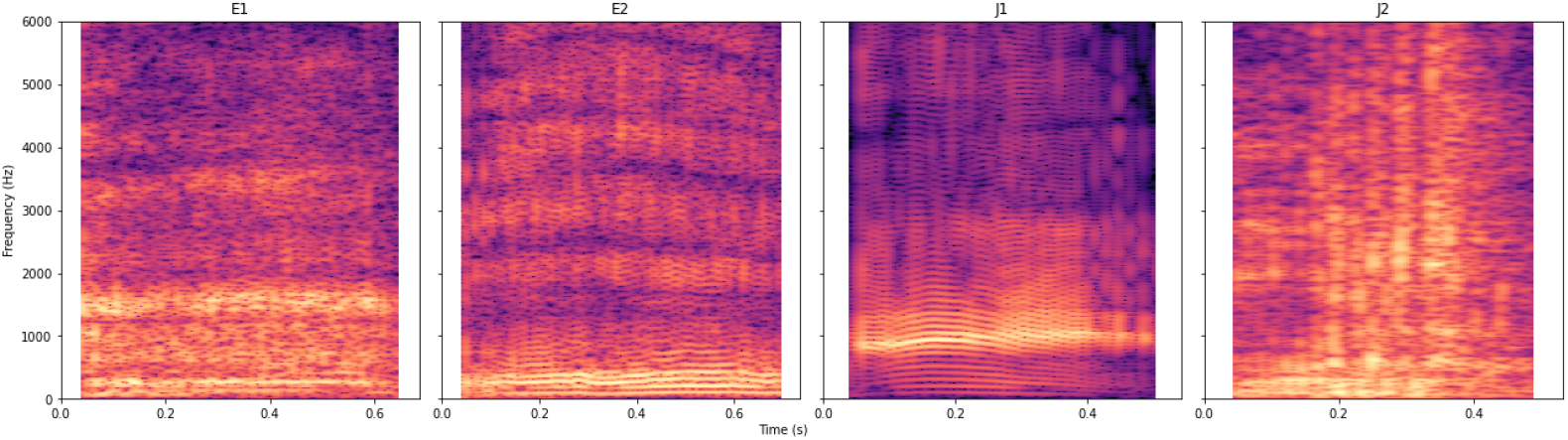
Spectral representation of the four vocalizations: E1, E2, J1 and J2. Frequency (Hz) on the y-axis, time (s) on the x-axis. Spectrogram was obtained with the *Parselmouth* package in Python (version 0.4.1, Praat 6.1.38; window length = 0.02 s, dynamic range = 70 dB, max. frequency = 7000 Hz (Boersma & Weenink, 2022; Jadoul et al., 2023a; Jadoul et al., 2018)).

We assumed that the seals would have formed a first association when they responded to approx. 50% of hand signs with the desired behavior. As a next step, we occasionally reinforced the seals for not vocalizing, i.e., for silence. We then trained the seals to associate this silence with a hand sign until they responded correctly to approx. 50% of hand signs (see Table 1). The association of these behaviors and their respective S_D_’s were interspersed in the seals’ daily research training sessions. During this stage, only the number of training sessions was noted, not the number of each requested behavior. This opportunistic training approach resulted from the circumstance that at this stage of the study, the seals could not be trained individually, and an assistant was not available.

#### Research Enclosure Training

In order to facilitate one-on-one testing sessions with the seals, a research enclosure was built. This enclosure was accessible through a gate to the main enclosure. Separating the seals and working with single individuals in the closed research area was trained through successive approximations to promote a gentle transition from group training to individual training. Initially, the seals were acquainted with the novel enclosure through occasional feeding at the gate. Incrementally, the seals were trained to follow into the area. We took small steps, so that after a short habituation phase and before the seals showed any signs of nervousness, they were sent into the pool (adjacent to the research enclosure). Sending into the pool served as a secondary reinforcement. The gate was kept open at all times during this stage. Once the seals’ stay in the research enclosure reached a couple of minutes without any signs of distress, closing of the gate was initiated. Here, one trainer rewarded the seal’s behavior (remaining rested), while another trainer closed and opened the gate in very small increments. After approximately five minutes of remaining rested without any signs of distress, we considered the seals comfortable with the research enclosure.

### Testing

All testing sessions occurred in the separate research enclosure. Trials alternated in a predetermined randomized order, which featured a counterbalanced number of trials and ≤ 3 trials of the same type in a row (Gellermann, 1933; Jadoul et al., 2023b). Once the seal was stationed, a trial began, and the trainer presented the respective hand sign (Fig. 2, Table 1).

In rare cases, when an animal was distracted (e.g., due to disturbance by zoo visitors) or vocalized incessantly, the trainer repeated the trial. The decision to repeat a trial was taken immediately by the trainer. Testing was conducted in the months of September and October 2021, and March and April 2022 (see Fig. 1).

#### Experiment 1: Vocalize - Silence

Sessions consisted of 20–50 trials/session, depending on the animal’s motivation; Sessions could be terminated when the animal lacked motivation, e.g., was distracted, chewed on fish rather than swallowing, left the experimental area, etc. Trials were counterbalanced between presenting the S_D_ for vocalizing and withholding from vocalizing, and presented in a pseudo-random order (Gellermann, 1933). The learning criterion was reached when the animal responded ≥ 80% correctly in four consecutive sessions.

#### Experiment 2: Produce Different Vocalizations Upon Different S_D_’s

Each session consisted of 20 trials. Trials were counterbalanced and interchanged between the different vocalization types, i.e., asking seal E for E1/E2 and seal J for J1/J2, and presented in a pseudo-random order (Gellermann, 1933). The learning criterion was reached when the animal responded ≥ 80% correctly in four consecutive sessions.

*Experiment 2* started when the seals had successfully completed *Vocalize vs. Silence* (Fig. 1). Before starting this experiment, seal E was trained on a second vocalization. This vocalization, E2, was established by selectively shaping the initial vocalization (E1) over a period of roughly four months, see section *Vocal Training*.

### Documentation

During testing, responses were noted on template response sheets (see Fig. 2). A subset of training and testing sessions were also audio and video recorded (Canon Legria HF25 camcorder). Audio recordings were obtained via a Zoom H6 digital recorder connected to a Sennheiser ME-67 unidirectional microphone (frequency response 40–20.000 Hz ± 2.5 dB) covered by a foam windshield.

### Data Analysis

We annotated audio recordings in Praat v6.1.38 (Boersma & Weenink, 2022). Relevant acoustic parameters of vocalizations were then batch-extracted with the *Parselmouth* package (a Python library for Praat, v0.4.1; Jadoul et al., 2023a; Jadoul et al., 2018) in Python. We chose acoustic parameters that were previously analyzed in harbor seal vocalizations, i.e., duration, spectral center of gravity, dominant frequency, percentage voiced, and median harmonicity (see Table 2; Raimondi et al., under review; Van Parijs et al., 2000).

**Table 2:** Definition of the acoustic parameters and description of the extraction procedure.

| Name of Acoustic Parameter | Description of Acoustic Parameter |
| --- | --- |
| Duration (s) | Length of the vocalization in time. |
| Spectral center of gravity (Hz) | The weighted average frequency of the vocalization's spectrum. The spectrum was calculated with Praat's "Sound: To Spectrum", default parameters. Then, the center of gravity was calculated using "Spectrum: Get center of gravity", default parameters. |
| Percentage voiced (%) | The pitch was calculated with Praat's "Sound: To Pitch" with default parameters, except pitch floor = 50 and pitch ceiling = 400). The number of voiced frames was divided by the number of all frames. |
| Median harmonicity (dB) | Harmonicity was calculated with Praat's "Sound: To Harmonicity (cc)", default parameters except minimum pitch = 50). After this the median of the harmonicity values were calculated. |
| Dominant frequency (Hz) | The spectrum was calculated with Praat's "Sound: To Spectrum" (default parameters). Then, we selected the frequency bin with the highest power. |

## Results

### Experiment 1: Vocalize - Silence

Seal E completed the first experiment within 5 sessions, i.e., 118 trials (23.6 trials/session ± 4.5) with 22 errors to criterion, the number of false trials until reaching the learning criterion. Seal J completed the first experiment within 6 sessions, i.e., 220 trials (36.6 trials/session ± 9.4) with 40 errors to criterion (see Figs. 4, 5, and 6). Before testing, Seal E had 9 preceding training sessions, and seal J had 15 preceding training sessions.

**Fig. 4:**
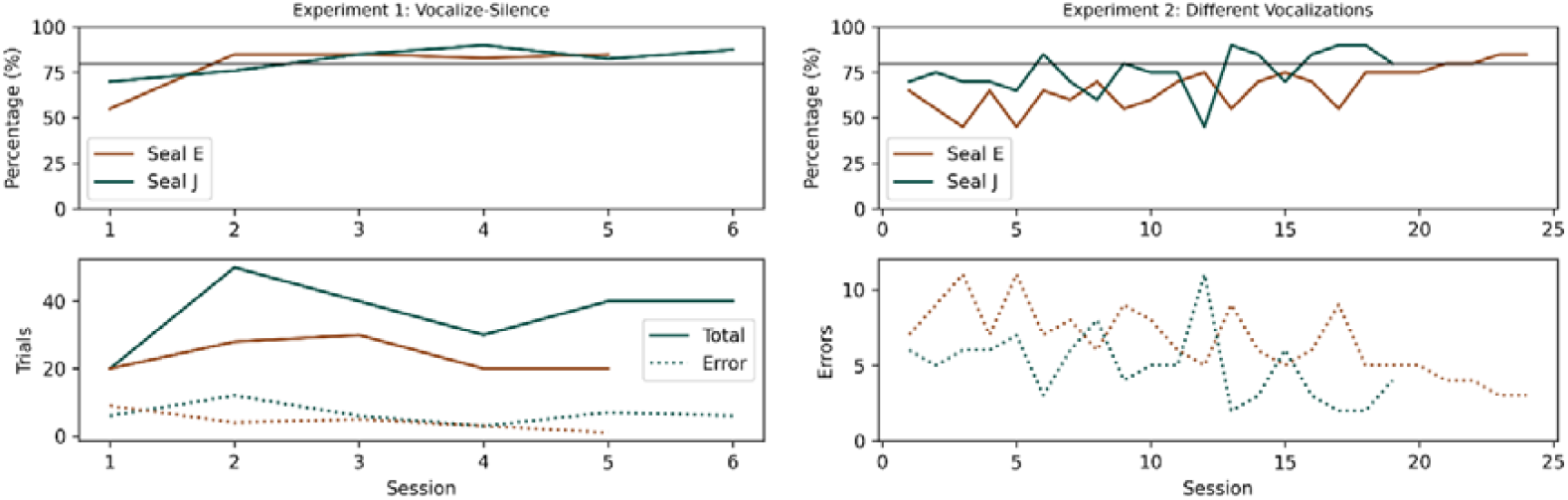
The top line plots depict the learning curves of seals E (brown) and J (green) for *Experiment 1* (top left) and *Experiment 2* (top right). The solid line at 80% on the y-axis marks the learning criterion’s threshold. The bottom line plots show the seals’ errors to criterion (dotted line), and total trial number (solid line) per session. Note that the total trial number per session varied for *Experiment 1* (see subsection *Experiment 1: Vocalize – Silence* in Material and Methods) but was consistent with 20 trials/session in *Experiment 2*.

**Fig. 5:**
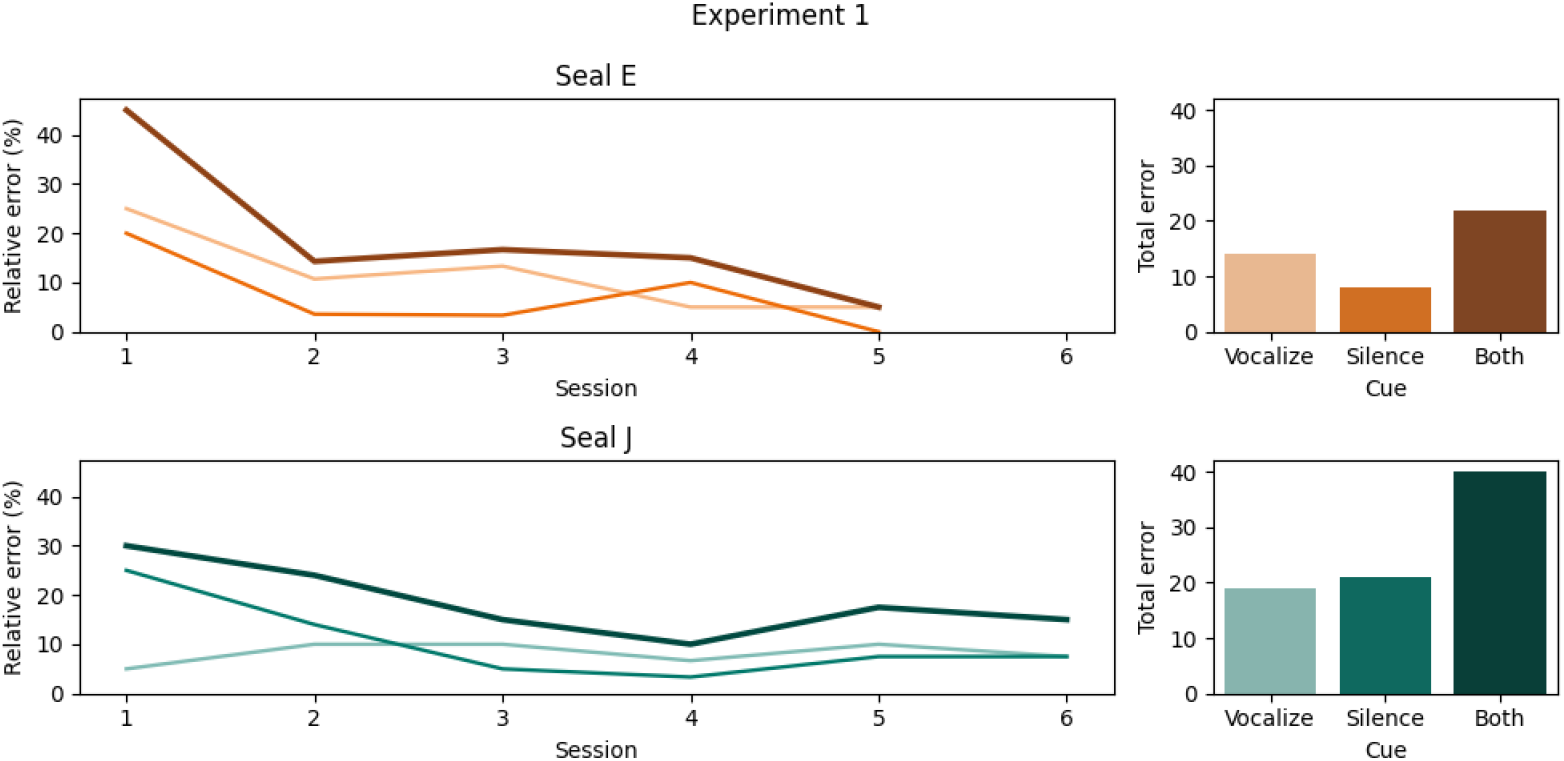
The line plots (left panel, seal E: top, seal J: bottom) depict the relative error of each session of *Experiment 1* for each S_D_ (*Vocalize, Silence*) separately, as well as the total relative error per session (dark brown/green). The bar plots (right panel, seal E: top, seal J: bottom) show the overall scores for each S_D_ over all sessions, as well as the total error to criterion.

**Fig. 6:**
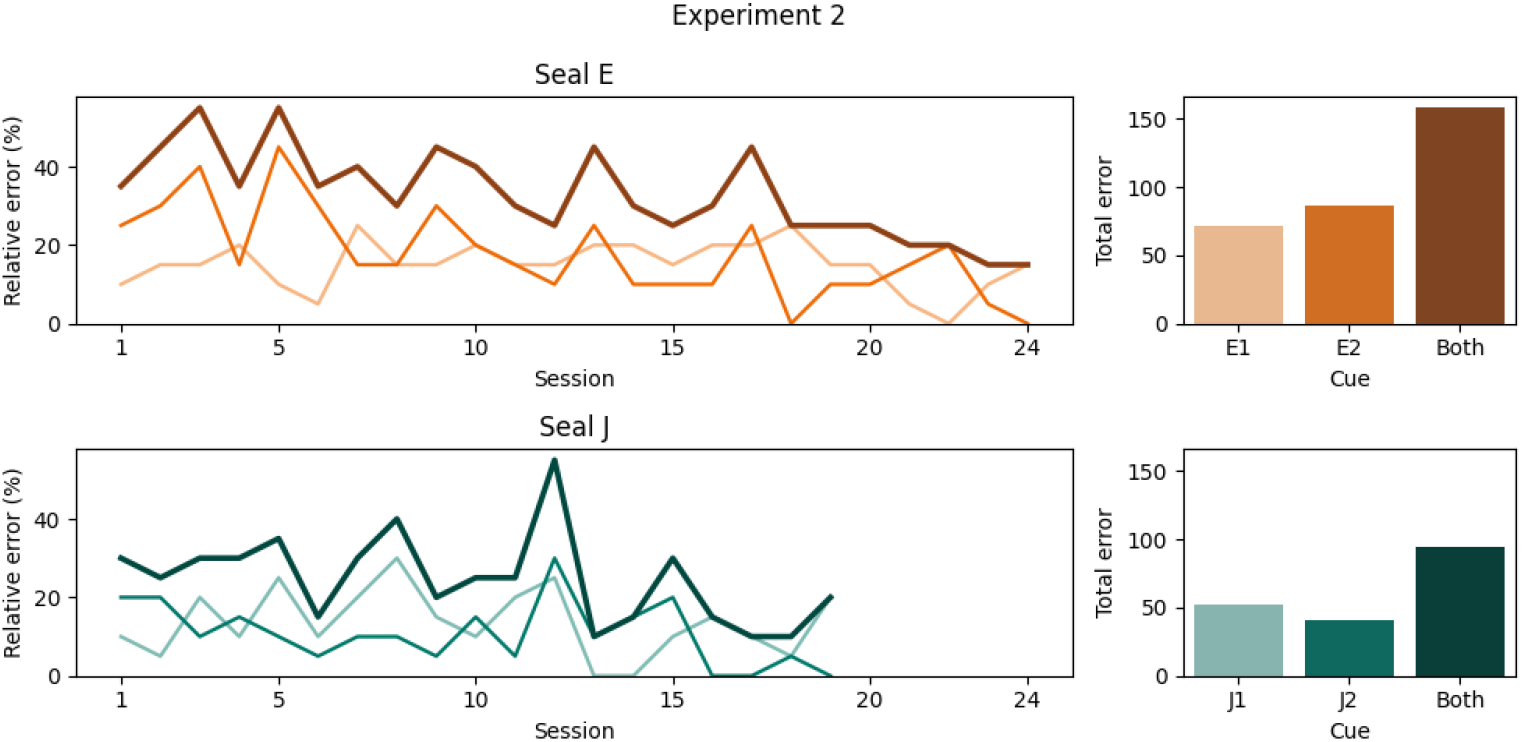
The line plots (left panel, seal E: top, seal J: bottom) depict the relative error of each session of *Experiment 2* for each SD (*E1, E2 and J1, J2*) separately, as well as the total relative error per session (dark brown/green). The bar plots (right panel, seal E: top, seal J: bottom) show the overall scores for each SD over all sessions, as well as the total error to criterion.

### Experiment 2: Produce Different Vocalizations Upon Different S_D_’s

Seal E completed the second experiment within 480 trials (24 sessions with 20 trials/session) with 158 errors to criterion. Seal J completed the second experiment within 380 trials (19 sessions with 20 trials/session) with 94 errors to criterion (see Figs. 4, 5, and 6). Four sessions were omitted from analysis as these were terminated early by seal J after a few trials. Before testing, Seal E had 47 preceding training sessions; this number includes the extra training she required to establish a novel vocalization type. Seal J had 23 preceding training sessions.

### Discernibility of Vocalization Types

To confirm that the seals produced different vocal types (E1 vs. E2 and J1 vs. J2) and that the types were not mere human subjective categories, we inspected the acoustic parameter composition between call types and within seals (Fig. 7).

**Fig. 7:**
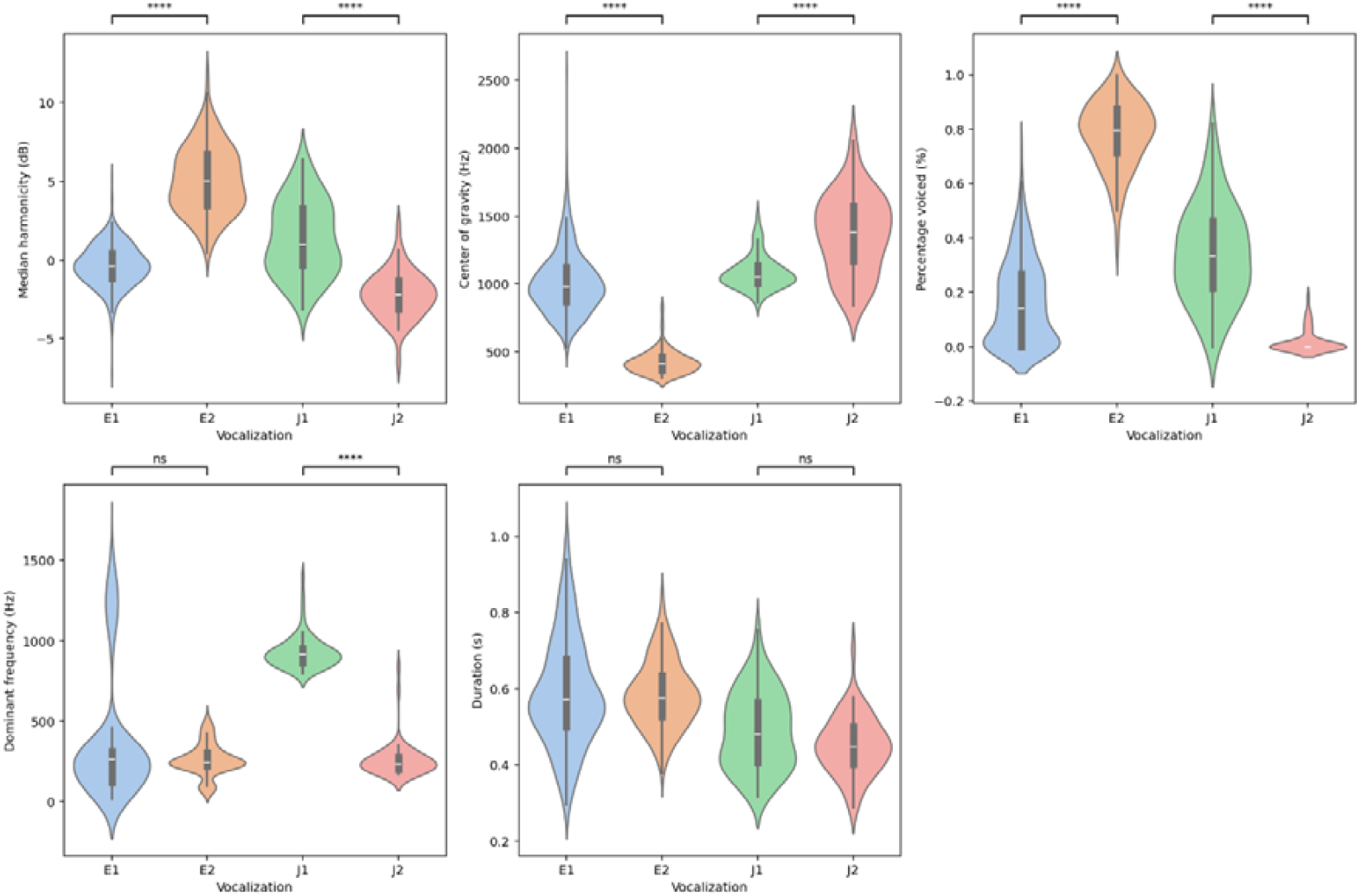
Visualized spectral parameter distribution of each seal’s vocalization types. Significance levels are indicated by asterisks, where ****: p < 0.0001, and ns: p > 0.05. The “median harmonicity” values are only shown in the range above -50 dB.

A Mann-Whitney-U test revealed that seal E’s two vocalization types differed significantly in three of the five extracted spectral parameters, namely median harmonicity (statistic = 457.0, p < 0.0001), center of gravity (statistic = 52608.0, p < 0.0001), and percentage voiced (statistic = 171.5, p < 0.0001). Seal J’s two vocalization types differed significantly in four of the five extracted spectral parameters, namely median harmonicity (statistic = 3825.0, p < 0.0001), center of gravity (statistic = 939.0, p < 0.0001), percentage voiced (statistic = 4349.5, p < 0.0001), and dominant frequency (statistic = 4470.0, p < 0.0001). P-values were Bonferroni-corrected for multiple comparison by a factor of five.

## Discussion

This study provides a protocol to train experimentally naive seals for a vocal learning experiment. In particular, we show how two harbor seals can be trained to associate different S_D_’s with different vocal behaviors, which demonstrates their ability of vocal usage learning.

The task in *Experiment 1*: *Vocalize vs. Silence*, was accomplished fairly fast, i.e., within 118 and 220 trials (seal E and J respectively). The task in *Experiment 2: Produce Different Vocalizations Upon Different S*_*D*_*’s*, was learned within 480 trials and 380 trials (seal E and J, respectively). A qualitative comparison between the average trial number of each experiment (*Vocalize vs. Silence*, x□ = 169 trials; *Produce Different Vocalizations Upon Different S*_*D*_*’s*, x□ = 430 trials) reveals that the seals needed over two and a half times more trials to learn to *Produce Different Vocalizations Upon Different S*_*D*_*’s*. Perhaps, this second experiment was more cognitively demanding than the first experiment; this difference would support the hypothesis that vocal usage learning features components of varying complexity (Janik & Slater, 2000; Shapiro et al., 2004; Vergara & Barrett-Lennard, 2017), which reflects in the time required to learn tasks.

During *Experiment 1*, the seals were asked to vocalize or remain silent on S_D_. Both seals succeeded at this task (Fig. 4). Investigating the error to criterion (ETC) value separated per S_D_ revealed that seal J responded almost equally correct (or incorrect) to either of the S_D_’s. Seal E, however, responded more often incorrectly to *Silence* than to *Vocalize* (Fig. 5). In *Experiment 2* the seals had to produce different vocalizations upon presentation of different S_D_’s. Here, seal E showed a higher ETC value for the E2 vocalization, than for the E1 vocalization (Fig. 6). A reason for this may be the active shaping of the E1 vocalization. This shaping directly preceded *Experiment 2*, causing a temporarily higher exposition of vocalization E1 over E2, which could have led to a type preference. Seal J showed similarly higher ETC values for one vocal type (J1) over the other (J2). It is not rare that animals show side or S_D_ preferences (Erdsack et al., 2022; Schluessel et al., 2014), which could be an underlying reason for the observed trends here.

In classical cognition tasks, considering the Clever Hans effect is important to avoid falsified results. The Clever Hans effect occurs, when an experimenter involuntary cues an animal. It leads to the animal learning a task based on other cues than intended (Sebeok & Rosenthal, 1981). Vocal usage learning, here, is independent of this phenomenon, as it is per definition learning to vocalize in association with a certain context (Janik & Slater, 2000; Janik & Slater, 1997). Whether or not the tested animals picked up on the intended S_D_’s or unintended cues is irrelevant. Vocal usage learning would be displayed in any case, solely on other cues than anticipated (Shapiro et al., 2004).

To study the learning process will require more fine-grained documentation of the training sessions from the moment of capturing a novel behavior, including trial numbers. This was unfortunately not possible in this study during training sessions. Further, a consistent number of trials per session enables the comparison of learning processes. Following such guidelines will further facilitate the direct comparison of vocal learning abilities in different individuals or species (Lattenkamp et al., 2021). Future studies should address this and track stimulus exposition during the training to better understand underlying (vocal) learning mechanisms.

Overall, our study shows that harbor seals, just like the closely related gray seal (Shapiro et al., 2004; Stansbury et al., 2015), can be trained to participate in vocal learning experiments. Through successive approximations they can be trained to remain rested in an external enclosure and there tested for the behaviors of interest. *Experiment 1* shows fast learning of differentiating vocalizing and remaining silent upon demonstration of different visual S_D_’s. *Experiment 2* shows that the seals are further capable of distinguishing two different vocalization types in response to different S_D_’s. Future studies should investigate whether harbor seals are also capable of vocal comprehension learning, i.e., reacting adequately to vocalizations in specific contexts.

## Conclusion

This study aimed at preparing two experimentally naive harbor seals for vocal learning experiments. Our report complements a previous one on vocal shaping and usage learning in a male harbor seal (Schusterman, 2008). Here, we show that harbor seals are good candidates for research, as they can be readily trained for participating scientific experiments. Further, this report confirms that male harbor seals are capable of vocal usage learning, and adds that female harbor seals are also capable of it. Our results further suggest that vocal learning may imply different levels of complexity, which to the best of our knowledge, has not been shown before in this species. Future studies should incorporate a tracking of the learning progress for a quantitative documentation of vocal usage learning, i.e., during the training phase all presented S_D_’s should be noted. It would be further interesting to build on Shapiro et al. (2004) and Stansbury et al. (2015) experimental design on higher levels of usage learning, to see whether harbor seals are capable of such “more complex” vocal usage learning.

## Acknowledgments

We thank Dr. Sara Torres Ortiz and PD Dr. Vera Schluessel for providing helpful comments on the manuscript. We thank Yannick Jadoul for helpful thoughts along the way, the discussions on stimulus randomization, and the brainstorming on plotting. Center for Music in the Brain was funded by the Danish National Research Foundation (DNRF117). The Comparative Bioacoustics Group was funded by Max Planck Group Leader funding to AR.

## Author Contributions

Conceptualization: DD, AR; Data Curation: DD; Formal Analysis: DD; Funding Acquisition: AR; Investigation: DD; Methodology: DD, AR; Project Administration: DD; Resources: DD, AR; Software: DD; Supervision: AR; Validation: DD; Visualization: DD; Writing – Original Draft Preparation: DD; Writing – Review & Editing: DD, AR

## Ethics Approval

Ethical review and approval were waived for this study, due to no permit requirement of the general training procedures that were conducted (Az. 81-04.78, Landesamt für Natur, Umwelt und Verbraucherschutz NRW).

## Consent for Publication

Not applicable.

## Data Availability

The data can be obtained from the corresponding author upon request.

